# NR0B1 alters the 9-*cis*-retinoic acid response in Ewing Sarcoma cells

**DOI:** 10.1101/2025.09.29.679070

**Authors:** Benjamin D. Sunkel, Meng Wang, Rachel D. Dreher, Melissa Sammons, Lindsay Ryan, Amy C. Gross, Scott Friedland, Ryan D. Roberts, Emily R. Theisen, Elaine R. Mardis, Richard K. Wilson

## Abstract

Pediatric cancers are often characterized by relatively low DNA mutation rates and are frequently driven instead by alterations in gene expression programs. In Ewing Sarcoma (EwS), these changes result from the activity of an oncogenic fusion transcription factor, EWS::FLI1, which alters the enhancer landscape and rewires transcriptional programs to promote oncogenic transformation. Among the targets directly activated by EWS::FLI1, NR0B1 has emerged as a potentially targetable transcription co-regulator and as a manifold for widespread alterations in downstream gene expression programs mediated by members of the ligand-inducible nuclear receptor superfamily of transcription factors. We have dissected the gene regulatory activity of NR0B1 in EwS models, showing its role in altering the basal gene expression programs of nuclear receptors COUP-TFII, EAR2, RXRa, and TR4. Additionally, we show that NR0B1 impacts the EwS response to retinoids, particularly 9-*cis*-retinoic acid signaling in part through RXRa. Our findings suggest NR0B1 silencing or inhibition offer a means of achieving a more potent response to retinoids, which may activate differentiation programs, decrease EwS gene expression signatures, and limit *in vitro* transformation phenotypes. Taken together, our study presents evidence of NR0B1 acting canonically as a nuclear receptor co-regulator in EwS, revealing a potential pathway to treatment of this aggressive cancer by combining NR0B1 inhibition with therapeutic nuclear receptor ligands.

## Introduction

Ewing Sarcoma (EwS) is an aggressive cancer occurring in bone and soft tissue throughout the body with peak incidence around 15 years or age [1]. A large number of patients present with micrometastases, informing the standard treatment regimen combining localized surgical resection and radiotherapy with systemic chemotherapy. Over decades of advancement and intensification of these therapies, 5-year survival rates for patients with local disease have improved markedly. Still, relapse occurs in 25% of patients (over 60% for those with metastatic disease), and survival rates following recurrence are dismal [2]. Making matters worse, EwS survivors exposed to aggressive radiation and intensified chemotherapy have an increased risk of secondary malignancies that is dose dependent [3, 4]. Addressing the urgent need for additional, effective treatment options for EwS patients, research teams have explored a range of immunotherapeutic strategies as well as tyrosine kinase and PARP inhibition, with several options advancing to clinical trial [5].

Compelling options for novel EwS treatments center on the disrupting the gene expression programs supporting disease growth. EwS is characterized by chromosomal translocations that produce an oncogenic fusion transcription factor, most often EWS::FLI1 [6]. EWS::FLI1 acts as a central node in directing EwS gene expression by co-opting chromatin regulatory protein complexes [7], altering the enhancer landscape [8], and driving changes in genome architecture [9, 10]. Efforts to target EWS::FLI1-driven gene expression have focused on inhibiting binding of the fusion to DNA (trabectedin) [11], leveraging transcriptional addiction (CDK7/9 inhibition) [12, 13], and reversing fusion-dependent chromatin repression (LSD1 inhibition) [14–16].

Another fruitful strategy may lie in the identification of EWS::FLI1 target genes that are necessary for EwS survival and can be inhibited pharmacologically. One such factor is NR0B1 (DAX1), an orphan nuclear receptor activated upon EWS::FLI1 binding to a GGAA microsatellite repeat near the *NR0B1* promoter [17–20]. Once expressed, NR0B1 reportedly co-regulates EWS::FLI1 target genes and has been shown to be necessary for *in vitro* transformation and *in vivo* xenograft tumor formation by EwS cells [17, 19, 21]. NR0B1 expression is typically limited to the adrenal gland, testis, and ovaries, and was first discovered as the causative gene in the development of X-linked adrenal hypoplasia congenita and accompanying hypogonadotropic hypogonadism [22, 23]. NR0B1 has an unusual structure among members of the nuclear receptor family of transcription factors, exhibiting an N-terminal array of LXXLL peptide motifs common to nuclear receptor co-regulators in lieu of a DNA binding domain [24]. This unique structure underlies the canonical function of NR0B1 in regulating the ligand-dependent gene regulatory activity of various nuclear receptors, demonstrated for SF-1 in the adrenal gland [25, 26]. In cancer models, aberrant expression of NR0B1 was shown to disrupt the activity of hormone nuclear receptors AR and PR in prostate and breast cancer cells, respectively [27], but to date, this activity has not been explored in EwS. Notably, a covalent inhibitor of NR0B1 has been reported [28], but a definitive role for NR0B1 in EwS is lacking and its suitability as a drug target remains unclear.

In the current study, we have profiled the genome-wide binding and identified disease-relevant target genes of NR0B1 in a panel of EwS cell lines and patient derived xenograft (PDX) tissues. We reveal broad interaction between NR0B1 and a variety of nuclear receptors, most notably RXRa. In agreement with its role as a nuclear receptor co-regulator, we find that NR0B1 silencing or inhibition allows for a more robust response to the pan-RAR/RXR ligand, 9-*cis*-retinoic acid. Following NR0B1 disruption, 9-*cis*-retinoic acid activates genes related to mesenchymal lineage tissue development and represses characteristic EwS gene expression signatures, leading to a reduction in *in vitro* transformation phenotypes. Our findings suggest that restoring nuclear receptor activity by NR0B1 inhibition may be a viable strategy for sensitizing EwS to therapeutic nuclear receptor ligands.

## Results

### Variable NR0B1 binding sites converge on common biological pathways in EwS cells

To gain a more comprehensive view of NR0B1 activity, we optimized the conditions for genome-wide NR0B1 ChIP-seq in four individual EwS cell lines: A673, SKNMC, TC32, and TC71. We found that NR0B1 binding sites, ranging in number from 18,961 in SKNMC cells to 80,917 in TC32 cells, were largely unique to individual cell types, with only 5,380 sites shared across all four EwS cell lines (Figure 1A). Despite this, NR0B1 binding site distributions with respect to gene coding regions were remarkably consistent across cell lines, and we noted more than 50% of NR0B1 sites occur outside of promoters and beyond 3 kb from an annotated transcription start site (TSS) (Supplementary Figure S1A,B).

**Figure 1.**
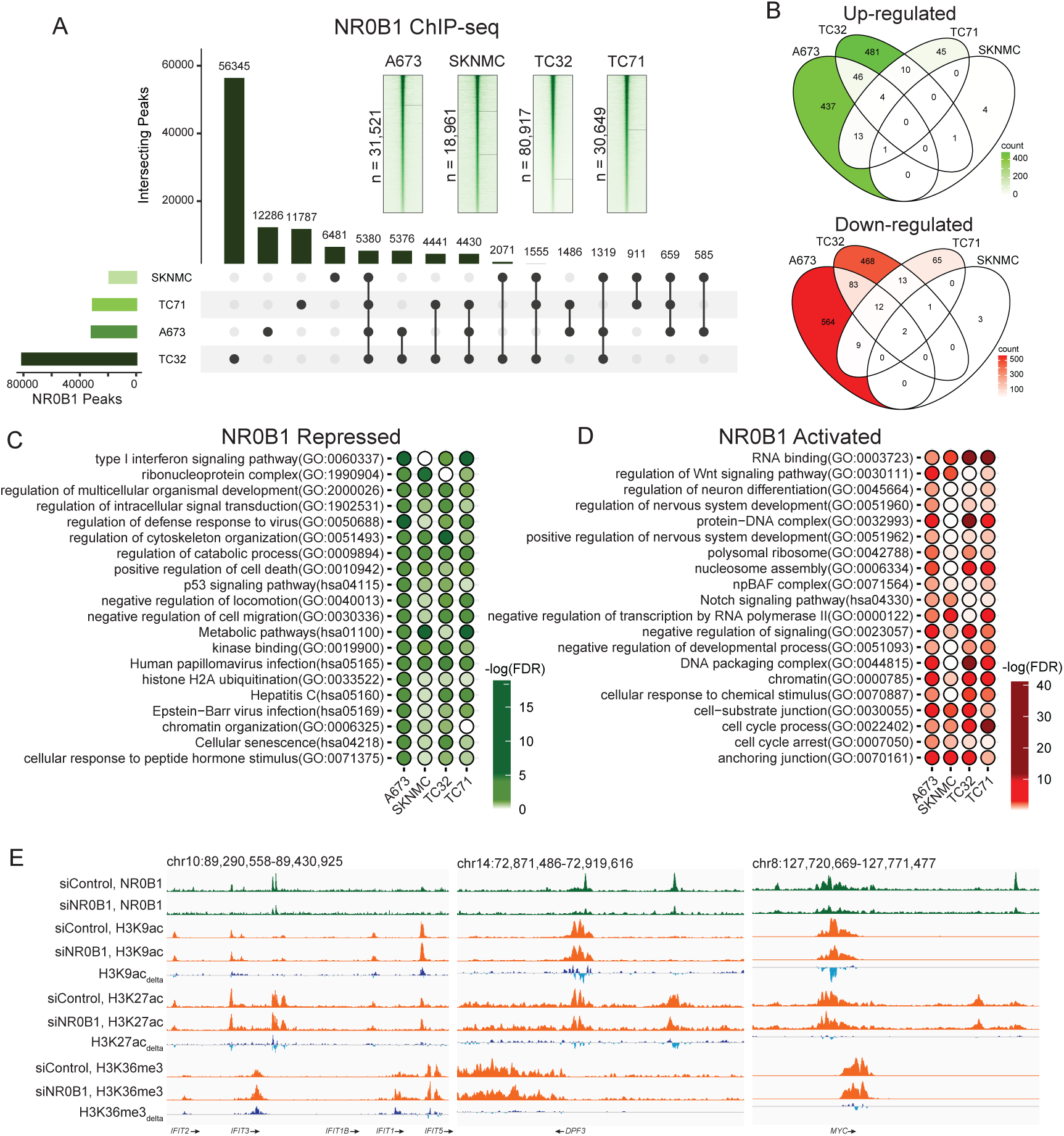
**A)** NR0B1 ChIP-seq was performed in A673, SKNMC, TC32, and TC71 EwS cell lines. Upset plot depicts shared and unique NR0B1 binding events across all cell lines. Inset heatmaps depict NR0B1 ChIP-seq signal strength across binding sites in each cell line. **B)** RNA-seq was performed in the indicated cell lines transfected for 72 hours with siControl or siNR0B1. Venn diagrams depict the common and unique differentially expressed genes (both up- and down-regulated) across cell lines resulting from NR0B1 silencing. **C-D)** NR0B1 binding sites and NR0B1-dependent gene expression profiles were combined on the Cistrome GO platform to predict putative NR0B1 target genes for gene ontology analysis. Balloon plots depict the pathways significantly enriched among NR0B1 repressed **(C)** and activated **(D)** targets in at least three of the four tested cell lines. **(E)** Integrative Genomics Viewer (IGV) plots of NR0B1, H3K9ac, H3K27ac, and H3K36me3 tracks in siControl and siNR0B1 transfected A673 cells NR0B1 repressed (*IFIT*) and activated (*DPF3* and *MYC*) loci. H3K9ac_delta_, H3K27ac_delta_, and H3K36me3_delta_ tracks represent the difference of (siNR0B1 – siControl) ChIP signals.

We next performed RNA-seq following transfection of each EwS cell line with Control-vs. NR0B1-targeting siRNAs (Supplementary Figure S1C). NR0B1 silencing resulted in highly variable gene expression changes across cell lines, where A673 cells showed 670 and 500 siNR0B1 up- and down-regulated genes, respectively, while SKNMC cells had only 12 genes significantly affected by loss of NR0B1 (Figure 1B). For integrated analysis of NR0B1 binding and NR0B1-dependent gene expression changes, we utilized the Cistrome-GO platform to derive regulatory potential scores for all putative NR0B1 target genes and compared the enrichment of these putative targets in Gene Ontology groups and KEGG pathways [29]. Despite heterogenous NR0B1 binding and impacts on gene expression, our Cistrome-GO analysis revealed highly convergent biological roles for putative NR0B1 target genes across the four EwS cell lines (Figure 1C,D and Supplementary Figure S1D). We consistently found that NR0B1 likely represses viral-induced innate immune pathways (e.g., interferon signaling), p53 and senescence pathways, response to hormone stimulation, and various catabolic processes while activating neuron differentiation pathways and the associated neural progenitor BAF (npBAF) chromatin remodeling complex, anchoring junctions, and cell cycle process (Figure 1C,D).

We asked if loss of NR0B1 effects the epigenome of EwS cells, underlying the gene expression changes we observed upon NR0B1 silencing. We performed ATAC-seq on siControl-vs. siNR0B1-transfected A673 cells, revealing 3,596 differential ATAC peaks upon NR0B1 loss (Supplementary Figure S1E). Combing these genomic regions with NR0B1-dependent gene expression changes, we compared putative targets of NR0B1 (Figure 1C,D) with putative targets regulated by our differential ATAC peaks and found a significant overlap between the two (hypergeometric overlap, corrected p-value = 8.4082×10^-4^) [30]. We noted that sites with reduced chromatin accessibility upon NR0B1 loss were often directly bound by NR0B1 (86.5%) and enriched for ETS DNA recognition motifs, indicating a direct role for NR0B1 in maintaining the accessibility of these sites, possibly in collaboration with EWS::FLI1 (Supplementary Figure S1E). On the other hand, 21.5% of sites with reduced accessibility were bound, albeit weakly, by NR0B1 and enriched with bZIP DNA sequence motifs (Supplementary Figure S1E). This indicates an indirect role for NR0B1 in compaction of these sites. We further profiled the chromatin modification landscape of A673 cells, confirming the localization of NR0B1 primarily to active enhancers and promoters by ChromHMM (Supplementary Figure S1F) [31]. Following NR0B1 silencing, we did not observe any global changes in the active or repressive histone modifications profiled across NR0B1 binding sites (Supplementary Figure S1G). Despite this global trend, we could readily identify individual gene loci where NR0B1 loss was associated with altered histone acetylation as well as gene body H3K36me3 levels reflective of differential gene expression, including the *IFIT* gene cluster on chromosome 10 and regulatory elements at the *MYC* and *DPF3* loci (Figure 1E). These data reveal that highly distinct NR0B1 binding sites across EwS cell lines function to regulate similar biological processes.

### NR0B1 regulates genes associated with poor EwS survival in vivo

To extend our findings of NR0B1 mediated gene expression, we performed bulk NR0B1 ChIP-seq as well as 10X single nuclei (sn)RNA/ATAC-seq on two EwS patient-derived xenograft (PDX) tissues, NCH-EWS-3 and NCH-EWS-5, which we confirmed express abundant NR0B1 protein (Supplementary Figure S2A). As in EwS cell lines, NR0B1 binding sites were mostly distinct across our two PDX tissues (Supplementary Figure S2B). We identified nuclei with high vs. low expression of NR0B1 in each PDX sample and performed differential gene expression and chromatin accessibility analyses between these two populations (Figure 2A,B and Supplementary Figure S2C). Similar to EwS cell lines, high NR0B1 expression correlated with activation or repression of largely unique gene sets between PDX tissues (Figure 2B). Yet, when we combined differential gene expression with NR0B1 binding site information, we found that putative NR0B1 targets from both tissues mapped to many of the same pathways (hypergeometric overlap, corrected p-value = 2.97×10^-13^). These included fatty acid metabolic programs, p53 signaling, hormone metabolism/signaling, and cell cycle/apoptotic processes.

**Figure 2.**
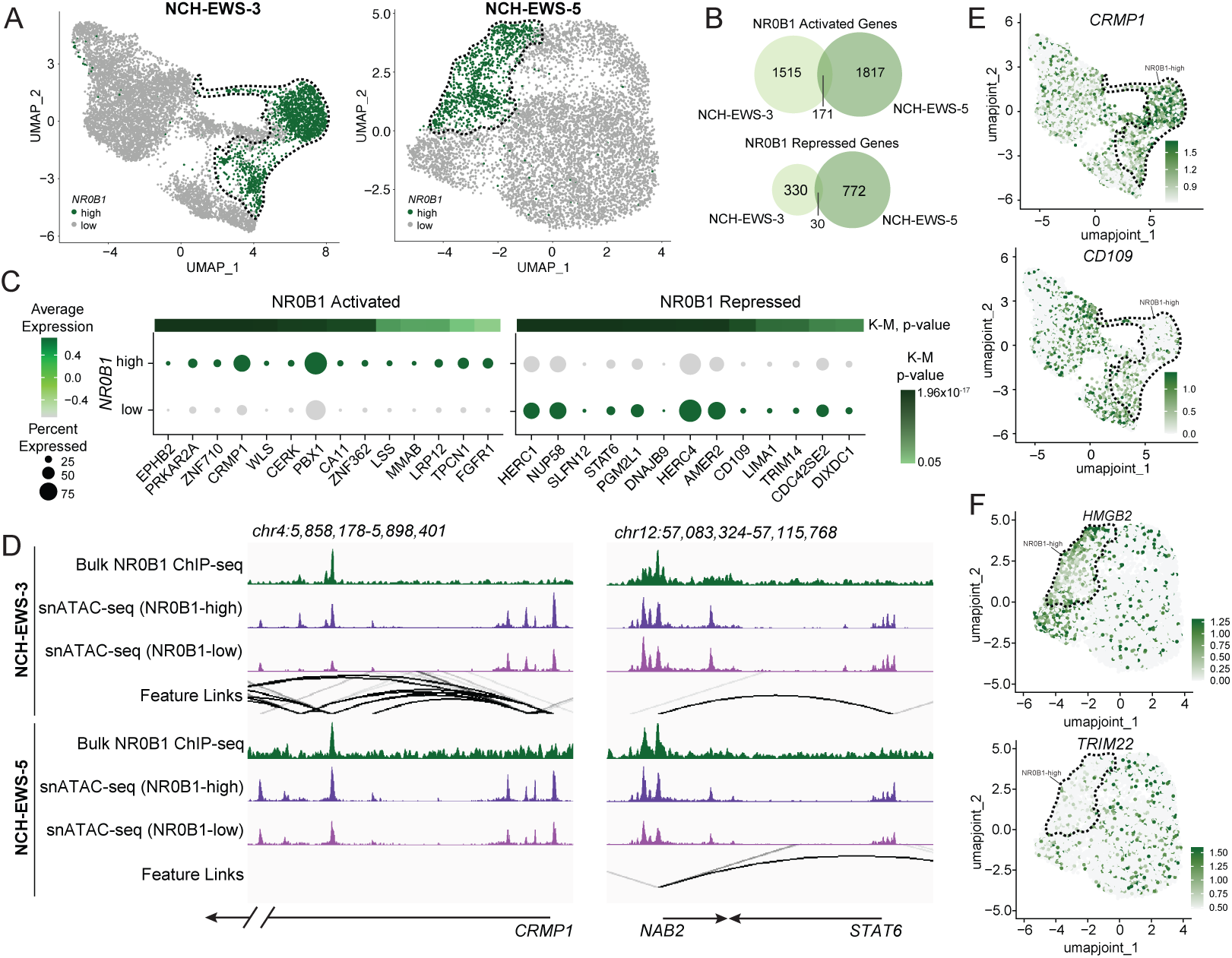
**A)** snRNA/ATAC-seq data from NCH-EWS-3 and NCH-EWS-5 PDX tissues were used to prepare joint ATAC/RNA UMAP projections and nuclei clusters are identified as expressing high vs. low levels of *NR0B1* (See Methods for details). **B)** Differential expression analysis was performed on *NR0B1* high vs. low clusters in each PDX. Venn diagrams depict the differential NR0B1 activated and repressed genes between NCH-EWS-3 and NCH-EWS-5. **C)** Genes activated or repressed by NR0B1 in NCH-EWS-3 as well as in A673 cells were uploaded to the R2 platform to conduct Kaplan-Meier survival analysis. Dot plot depicts the expression patterns (in NR0B1 high vs. low nuclei) of NR0B1 activated genes for which high expression in EwS patients is significantly associated with poor overall survival (left) and NR0B1 repressed genes for which low expression is associated with poor overall survival (right). Adjusted p-values for Kaplan-Meier analysis are provided for each gene. **D)** IGV plots display the binding of NR0B1 (bulk ChIP-seq), snATAC-seq signal (NR0B1 high vs. low nuclei), and feature links in each PDX tissue at the NR0B1 activated *CRMP1* locus and the NR0B1 repressed *STAT6* locus. **E-F)** Feature plots display the expression of NR0B1 activated (*CRMP1* and *HMGB2*) and repressed (*CD109* and *TRIM22*) genes in NR0B1 high vs. low nuclei in NCH-EWS-3 **(E)** and NCH-EWS-5 **(F)**.

We identified just 46 differential ATAC peaks in NCH-EWS-5 between NR0B1-high vs. - low nuclei, so we excluded this tissue from further analyses related to chromatin accessibility. In NCH-EWS-3, we found 3,720 peaks with enhanced accessibility and 1,350 peaks with reduced accessibility in the NR0B1-high population (Supplementary Figure S2C,D). Paralleling our cell line findings, sites with enhanced accessibility in the NR0B1-high population showed stronger binding of NR0B1 and enrichment of both ETS and bHLH motifs, suggesting a direct role for NR0B1 in maintaining chromatin accessibility of these sites (Supplementary Figure S2D). In regions of reduced chromatin accessibility in the NR0B1-high population, we noted low binding of NR0B1 and enrichment of both bZIP and ETS motifs. With an abundance of snATAC-seq signal changes between NR0B1-high and -low populations in NCH-EWS-3, we could rely on the feature linkage analysis in Seurat to highlight distal regulatory regions that may control individual gene expression changes. We found that the expression of *NR0B1* was correlated with differences in chromatin accessibility of a pair of EWS::FLI1-bound sites 81 kb upstream of the *NR0B1* locus. It may be of interest to consider these sites as important nodes in the network of regulatory elements driving *NR0B1* expression in addition to the previously reported proximal GGAA microsatellite repeat [19, 20].

We next winnowed our lists of putative NR0B1 targets in NCH-EWS-3 and NCH-EWS-5 by retaining only genes that showed concordant NR0B1-dependent gene expression in A673 cells. We utilized the remaining genes for Kaplan-Meier analysis in a cohort on EwS patients for which gene expression microarray and long-term survival data were available on the R2: Genomics Analysis and Visualization Platform [32, 33]. Through this effort, we identified 14 and 31 NR0B1 activated genes whose high expression is associated with shorter overall survival in EwS (in NCH-EWS-3 and NCH-EWS-5, respectively), as well as 13 and 5 NR0B1 repressed genes for which low expression is associated with shorter overall survival (in NCH-EWS-3 and NCH-EWS-5, respectively) (Figure 2C and Supplementary Figures S2F,G). Looking at individual gene loci, we find that enhanced expression of *CRMP1* in NR0B1-high nuclei is associated with enhanced accessibility of an intronic site that is occupied by NR0B1 in both PDX tissues (Figure 2D). Similarly, reduced expression of *STAT6* in the NR0B1-high population is correlated with chromatin accessibility changes at a region just upstream of the *NAB2* locus where we find evidence of NR0B1 binding in both PDX models (Figure 2D). We have highlighted the relative expression of *CRMP1* and *CD109* in NCH-EWS-3 as well as *HMGB2* and *TRIM22* in NCH-EWS-5 to demonstrate their respective activation and repression in

NR0B1-high nuclei in these tissues (Figure 2E,F). We find it noteworthy that *CD109* is the only NR0B1 target we identified across both PDX tissues, highlighting what appears to be a consistent repression of inflammatory or innate immune processes by NR0B1 across model systems that is linked to poor EwS outcomes (Figures 1C & 2C) [34].

### NR0B1 exhibits extensive interplay with nuclear receptors in EwS cells

We performed motif analysis of all NR0B1 binding sites identified in EwS cell lines and PDX samples. Despite occupying distinct genomic regions across the models we tested (Figure 1A and Supplementary Figure S2B), NR0B1 sites showed highly consistent enrichment of DNA sequence motifs (Figure 3A). We noted significant enrichment of ETS motifs recognized by EWS::FLI1 in all models, consistent with previous reports of NR0B1 colocalization with this EwS-driving fusion transcription factor [21]. Interestingly, we also identified a broad array of nuclear receptor (NR) motifs in NR0B1 sites from each model, reflecting a canonical role for NR0B1 as an NR co-regulator [24]. To follow up on this potential interaction between NR0B1 and other NRs expressed in EwS cells, we performed ChIP-seq for COUP-TFII (NR2F2), EAR2 (NR2F6), RXRa (NR2B1), and TR4 (NR2C2) in A673 cells. We found a strong degree of overlap between NR0B1 and each NR, ranging from 49.9% in COUP-TFII sites to 67.5% in RXRa sites (Figure 3B). These overlaps exceeded that observed for EWS::FLI1 with NR0B1 (20.2%) [35]. In most cases, NR0B1/NR overlapping sites were found in active promoters and enhancers (Supplementary Figure S3A). Noting the relatively strong colocalization between NR0B1 and RXRa, we confirmed this observation in additional EwS cell lines and found overlaps ranging from 37.3% in SKNMC cells to 71.3% in TC32 cells (Supplementary Figure S3B).

**Figure 3.**
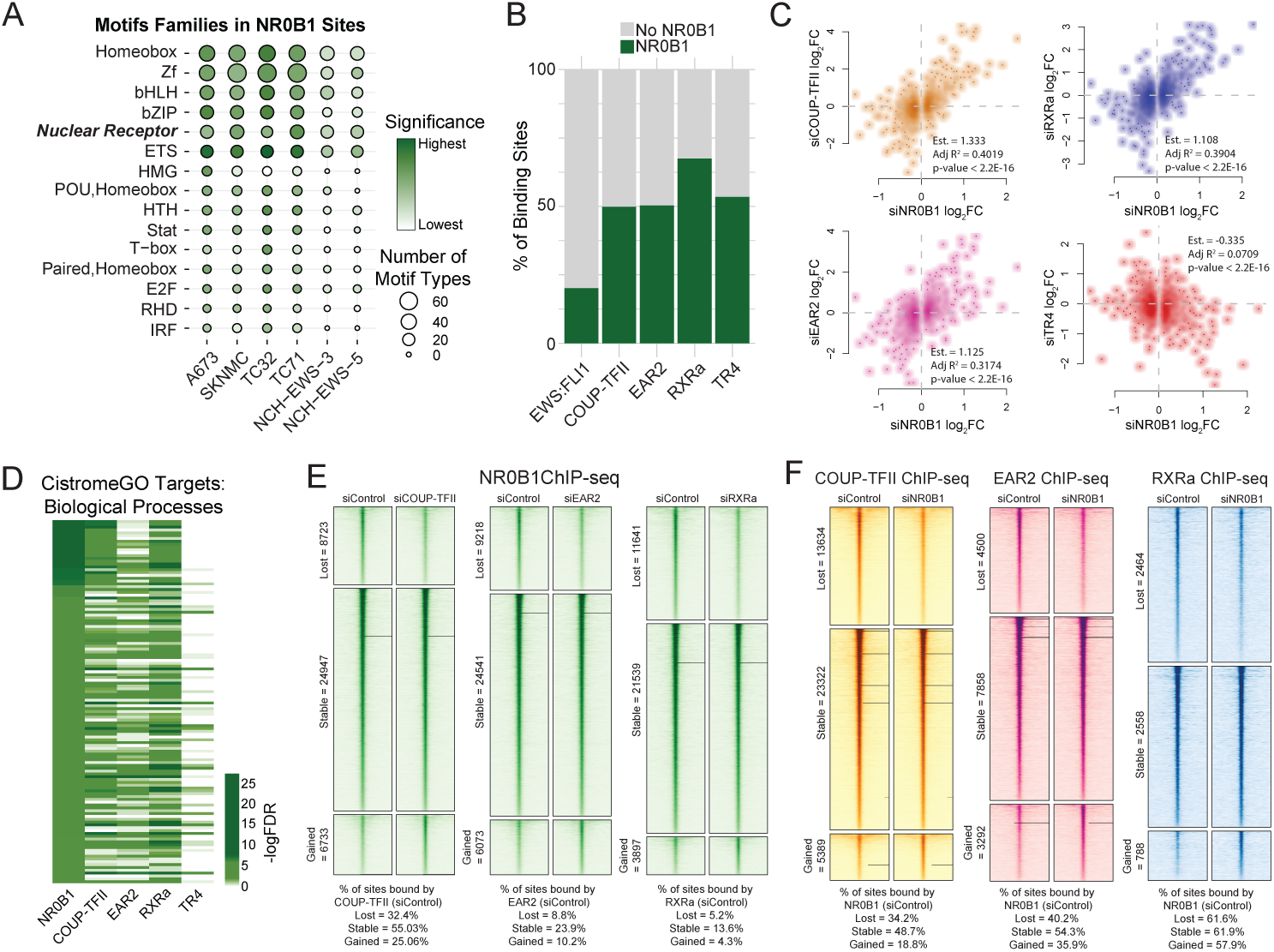
**A)** Motif analysis was performed on NR0B1 binding sites detected in the indicated EwS cell lines and PDX models. Balloon plot depicts the number of individual motif types within each motif family, and the relative significance of the enrichment for each motif family across all cells/tissues. **B)** Overlap of NR0B1 binding with sites occupied by EWS::FLI1 or the NRs COUP-TFII, EAR2, RXRa, and TR4 in A673 cells. **C)** RNA-seq was performed in A673 cells transfected with siControl or siRNA targeting COUP-TFII, EAR2, RXRa, or TR4. Scatter plots display the fold change (log_2_FC) in gene expression resulting from NR silencing vs. NR0B1 silencing. Inset reports the coefficient/estimate, adjusted R^2^, and p-value of linear regression models assessing the correlation between NR0B1 and NR-dependent gene expression changes. **D)** NR binding sites and NR-dependent gene expression profiles were uploaded to Cistrome GO to identify NR target genes and perform gene ontology analysis. Heatmap displays the significance of enrichment of NR0B1 regulated biological processes among NR target genes. **E)** NR0B1 ChIP-seq was performed following COUP-TFII, EAR2, or RXRa silencing in A673 cells. Heatmaps display the ChIP-seq signal and percent NR overlap in NR0B1 binding sites that are lost, stable, or gained as a result of NR silencing. **F)** ChIP-seq was performed for the indicated NR in siControl vs. siNR0B1 transfected A673 cells. Heatmaps display the ChIP-seq signal and percent NR0B1 overlap in NR binding sites that are lost, stable, or gained as a result of NR0B1 silencing.

We next asked if the colocalization of NR0B1 with other NRs in EwS cells resulted in shared regulation of target gene expression. We silenced individual NRs before performing RNA-seq, and we found that in general, NRs are expressed independent of one another (Supplementary Figure S3C). The one exception to this was the significant up-regulation of *NR0B1* upon silencing of COUP-TFII, EAR2, RXRa, or TR4. We found that gene expression changes resulting from NR0B1 silencing were generally recapitulated upon silencing of COUP-TFII, EAR2, and RXRa, while TR4-dependent gene expression changes were anticorrelated with NR0B1-driven changes (Figure 3C). When we performed Cistrome-GO analysis combining NR binding sites with NR-dependent gene expression, we found that predicted target genes of NR0B1 largely mapped to the same pathways as the predicted targets of COUP-TFII, EAR2, and RXRa (Figure 3D), which was not the case for TR4 target genes.

As NR0B1 lacks a DNA-binding domain (DBD), it is reportedly recruited to chromatin through interactions between its own protein domains (both its N-terminal array of LXXLL motifs and its C-terminal ligand-binding domain [LBD]) and those of the NRs it coregulates [35–37]. It follows that NR chromatin binding may determine NR0B1 genomic distribution patterns. To further characterize the interplay between NR0B1 and NRs in EwS, we assessed their hierarchical binding in A673 cells. We began by silencing COUP-TFII, EAR2, or RXRa and performing ChIP-seq against NR0B1. This revealed many thousands of NR0B1 sites impacted by NR loss, most often resulting in weakened/lost NR0B1 binding (Figure 3E). However, we noted that a relatively small number of NR0B1 sites lost upon NR silencing were occupied by the corresponding NR (ranging from 32.4% for COUP-TFII down to 5.2% for RXRa). This indicates that most of the effect of NR silencing on NR0B1 binding may be indirect. Similarly, sites with enhanced/gained NR0B1 binding upon NR silencing were infrequently bound by the corresponding NR, though there was evidence in a small fraction of sites that NRs may directly preclude NR0B1 occupancy (Figure 3E). Due to the up-regulation of *NR0B1* upon silencing COUP-TFII, EAR2, and RXRa (Supplementary Figure S3C), it is possible that a portion of gained NR0B1 binding sites may reflect additional NR0B1 in the cell.

Interestingly, when we silenced NR0B1, we also identified thousands of differential NR binding events, including the loss of nearly half of RXRa binding sites (Figure 3F). Relatively large percentages of these NR0B1-dependent NR binding sites were occupied by NR0B1, indicating that NR0B1 frequently stabilizes or prevents NR binding directly. We were intrigued by this result and looked to our epigenomic data for clues as to why NR0B1 loss could result in stronger or weaker binding of RXRa. While we observed modest alterations in H3K4 methylation status, these changes were observed in all three RXRa binding site groups (i.e., siNR0B1 lost, stable, and gained), and so they are unlikely to specifically explain enhanced or weakened RXRa binding upon NR0B1 silencing. Motif analysis of RXRa sites lost or gained after NR0B1 silencing revealed a variety of zinc finger, ETS, bZIP, and NR motif types, indicating relocalization to a broad array of potential binding partners. Taken together, these data provide evidence of widespread colocalization of NR0B1 with several NRs in EwS cells. This colocalization is associated with mutual coregulation of shared NR/NR0B1 target genes (for COUP-TFII, EAR2, and RXRa) as well as transcriptional antagonism (for TR4). Additionally, our data suggest a complex hierarchy between NR0B1 and NR binding patterns in which there is bidirectional influence leading to both stabilization and destabilization of binding by both direct and indirect means.

### NR0B1 alters the response of EwS cells to retinoids

Due to their strong genome-wide colocalization, mutual target gene regulation, and interplay in chromatin binding, we went on to ask if NR0B1 expression affects the ligand-responsive gene regulatory activity of RXRa. We performed RNA-seq in siControl and siNR0B1 transfected A673 and SKNMC cells treated with vehicle or 100 nM 9-*cis*-retinoic acid (9cRA, a pan-RAR/RXR agonist) for 24 hours [38]. In the presence or absence of NR0B1, 9cRA treatment resulted in roughly 1,800 and 1,300 differentially expressed genes in A673 and SKNMC cells, respectively. This included shared up-regulation of retinoid inducible fibroblast activation protein (*FAP),* retinaldehyde reductase *DHRS3*, and retinoic acid receptors *RARB* and *RARG* (Supplementary Figure S4A) [39–41], indicating that EwS cells can mount a retinoid response independent of NR0B1 status. However, we found that hundreds of 9cRA-responsive genes were uniquely altered only in siControl or siNR0B1 transfected cells (Figure 4A). Specifically, NR0B1 silencing resulted in 9cRA-dependent activation of interferon response and complement pathways as well as differentiation programs such as adipogenesis and myogenesis (Figure 4B).

**Figure 4.**
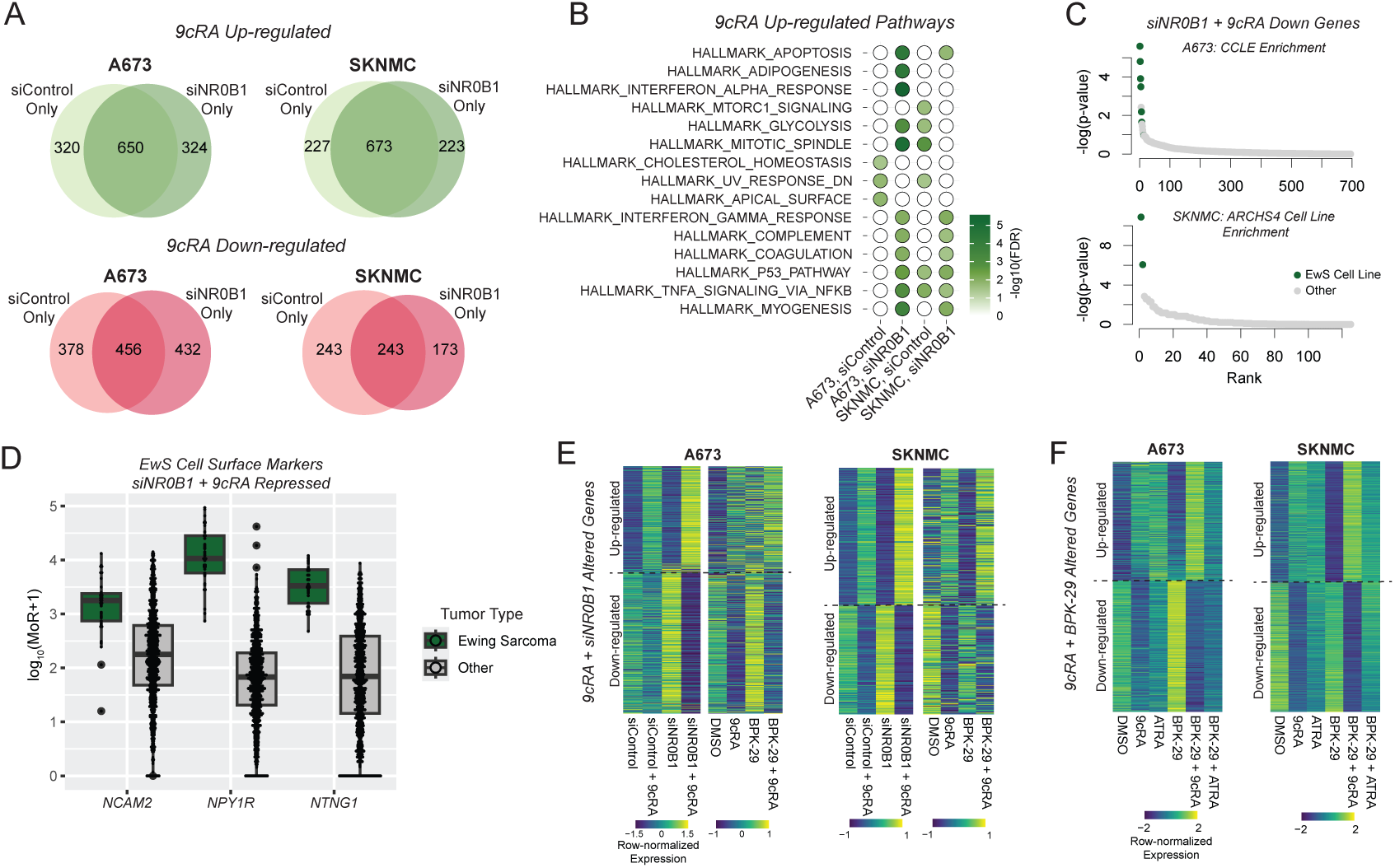
**A)** RNA-seq was performed in A673 and SKNMC cells treated for 24 hours with 100 nM 9cRA of Vehicle (DMSO). Differential expression analysis identified 9cRA responsive gene expression changes in siControl vs. siNR0B1 transfected cells. Venn diagrams display the genes uniquely up-regulated or down-regulated by 9cRA before and after NR0B1 silencing. **B)** Genes uniquely activated by 9cRA in siControl vs. siNR0B1 transfected cells were used to assess enrichment of MSigDB Hallmark Pathways. Balloon plot depicts pathway enrichment in siControl vs. siNR0V1 transfected cells. **C)** EnrichR analysis was performed to identify cell types characterized by genes specifically down-regulated by 9cRA in siNR0B1 transfected cells. Figure displays the cell line ranks (EwS cells or all others) and significance of enrichment for genes identified in A673 (top, CCLE annotation) and SKNMC (bottom, ARCHS4 annotation) cells. **D)** Expression level of the indicated cell surface marker genes in EwS vs. all other solid tumors. **E)** Comparison of gene expression changes driven by 9cRA treatment following NR0B1 silencing (siNR0B1) vs. inhibition (BPK-29) in A673 and SKNMC cells. **F)** Comparison of gene expression changes driven by 9cRA vs. ATRA in A673 and SKNMC cells treated with Vehicle (DMSO) or BPK-29.

Among genes down-regulated by 9cRA treatment, we found more similar enrichment of hallmark pathways between siControl and siNR0B1 transfected cells (Figure 4B). Using EnrichR [42], however, we observed that genes repressed by 9cRA only after NR0B1 silencing were characteristic of EwS cell lines (Figure 4C) [43, 44], indicating that 9cRA may drive a loss of Ewing identity when combined with NR0B1 disruption. Interestingly, we found multiple cell surface receptor and adhesion molecules among these genes, including *NTNG1*, *NPY1R*, and *NCAM2* (Supplementary Figure S4C). These genes exhibit high expression in EwS as compared to other pediatric solid tumors (Figure 4D), further reinforcing that 9cRA treatment in the absence of NR0B1 can inhibit the expression of characteristic EwS markers.

We were curious to find if NR0B1 inhibition with the recently reported covalent inhibitor, BPK-29 [28], would phenocopy NR0B1 silencing with respect to its impacts on the retinoid response. Interestingly, 24-hour treatment of A673 and SKNMC cells with 100 nM BPK-29 alone had minimal impact on their gene expression profiles. In A673, this was particularly notable, given the large number of genes altered by NR0B1 silencing (Figure 1B and Supplementary Figure S4C). Despite these differences, we found the 9cRA response was similarly impacted by NR0B1 silencing or inhibition (Figure 4E and Supplementary Figure S4D). We also asked if a second retinoid, all-*trans*-retinoic acid (ATRA), which is a ligand for RARs but lacks strong affinity for RXRs [38], has similar impacts on EwS gene expression when combined with NR0B1 inhibition. Though we did observe BPK-29-dependent changes in the response to ATRA (Supplementary Figure S4E), we were intrigued to find differences in the response to 9cRA vs. ATRA in the presence of BPK-29, where 9cRA resulted in more and stronger gene expression changes altering distinct pathways compared to ATRA (Figure 4F and Supplementary Figure S4F).

To begin to understand how BPK-29 alters the response to 9cRA in EwS cells, we performed RXRa ChIP following 9cRA treatment alone or in combination with BPK-29. When we specifically looked for RXRa binding sites stabilized by treatment with 9cRA, we noted large discrepancies between BPK-29-treated vs. untreated cells (Supplementary Figure S4G), including 3,661 9cRA-stimulated RXRa sites only detected in the presence of BPK-29. NR0B1 binding itself appeared to be unaffected by 9cRA or BPK-29 across these regions. Combining these distinct RXRa binding sites with genes altered by 9cRA alone or in combination with BPK-29, we found that the predicted targets of RXRa were largely unique when comparing 9cRA treatment with or without BPK-29 (Supplementary Figure S4H). Motif analysis identified the composite RAR/RXR motif to be highly enriched in each RXRa binding site category (Supplementary Figure S4I), indicating that NR0B1 action likely influences the distribution of RXRa to canonical retinoid response elements rather than directing RXRa to genomic regions associated with other transcription factors. Together, these data demonstrate a substantial alteration of the retinoid response in EwS cells resulting from NR0B1 expression and activity. We have shown that silencing or inhibition of NR0B1 paves the way for a more robust response to retinoids, particularly the pan-RAR/RXR agonist 9cRA. The result is partial activation of differentiation pathways and repression of genes uniquely characterizing EwS cells and tissues.

### BPK-29 enhances the phenotypic response to 9-cis-retinoic acid in EwS cells

Based on our observation that NR0B1 impacts the transcriptional response of EwS to 9cRA treatment, we asked if NR0B1 inhibition with BPK-29 would combine with 9cRA in altering EwS *in vitro* cell growth properties. In 2-dimensional cell proliferation assays, we found that 9cRA alone had little effect, while BPK-29 inhibited EwS cell growth with an IC_50_ around 1 μM in A673 cells and 650 nM in TC32 cells (Supplementary Figure S5A). Interestingly, A673 growth was stimulated across a broad range of 9cRA + BPK-29 dosing combinations, revealing partial antagonism between these compounds with respect to proliferation as a monolayer that was recapitulated in TC32 cells (Figure 5A and Supplementary Figure S5B). To assess the transformation phenotype of EwS cells exposed to 9cRA in combination with BPK-29, we utilized the Growth in Low Attachment (GILA) assay [45], in which EwS cells readily form spheroids, to screen for potential compound interactions. We found that 9cRA (at 5-10 nM) exhibited synergistic disruption of spheroid growth when combined with BPK-29 (at 10-250 nM) in A673 cells (Figure 5B). Similar results were seen in SKNMC and TC32 cells (Supplementary Figure S5C). To confirm these results, we assessed the anchorage-independent growth of A673 cells using the soft agar colony formation assay. We found that 100 nM BPK-29 alone reduced colony formation, while combining BPK-29 with 10 nM 9cRA further reduced colony formation (Figure 5C). We also noted trends in the reduction of A673 colony size when comparing dual BPK-29 + 9cRA treatments to single agents (Supplementary Figure S5D). Together, these early data link our observations of a molecular interaction between RXRa and NR0B1 to a phenotypic outcome in which NR0B1 disruption enhances the effect of 9cRA on the growth of EwS cells. Our GILA and soft agar colony formation results suggest some potential for NR0B1 inhibition to combine with RXR/RAR agonism to inhibit the oncogenic transformation of EwS cells.

**Figure 5.**
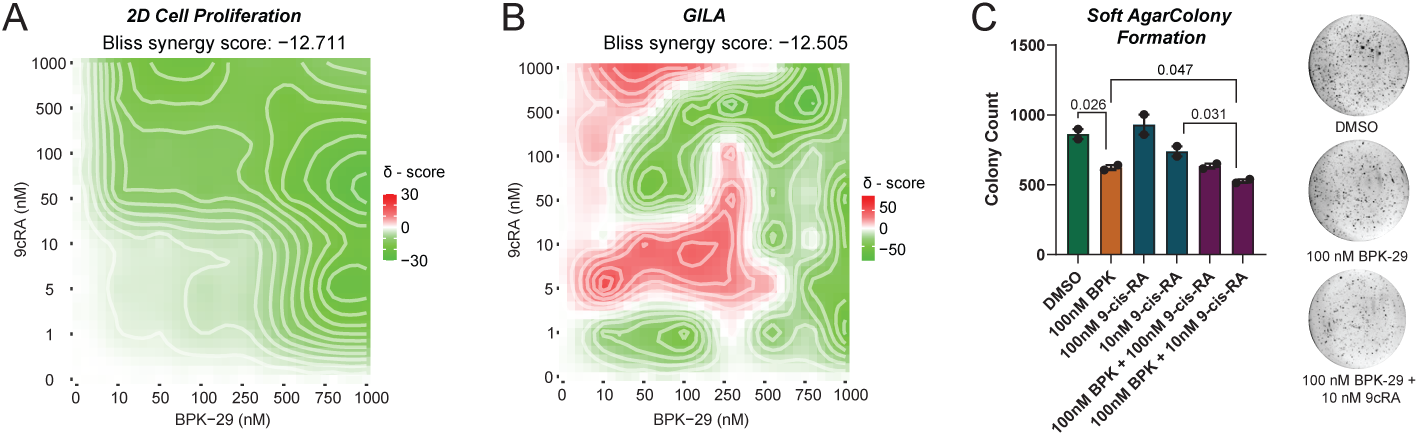
**A)** A673 cell grown in a monolayer were treated with the indicated combinations of 9cRA and BPK-29 for 6 days, and cell viability was measured. Results were analyzed on the SynergyFinder platform, demonstrating antagonism between 9cRA and BPK-29. **B)** A673 spheroids were grown in ultra-low attachment plates exposed to the indicated combinations of 9cRA and BPK-29 for 6 days. Viability results were analyzed on the SynergyFinder platform, demonstrating compound interaction between 9cRA and BPK-29. **C)** Soft agar colony formation assays were performed on A673 cells exposed to the indicated treatments. (Left) Colony counts are provided as well as p-values (t-test) for significant comparisons. (Right) Representative images of colonies formation with DMSO, 100 nM BPK-29, or 100 nM BPK-29 + 10 nM 9cRA.

## Discussion

These findings represent a significant advancement in our understanding of disease-relevant transcriptional mechanisms underlying EwS. While previous studies conclusively demonstrated direct activation of NR0B1 by EWS::FLI1 in EwS cells [17, 18], there is relatively little known about the subsequent function of this non-canonical orphan nuclear receptor in this disease setting. NR0B1 is reported to serve as a co-regulator of EWS::FLI1 and is deemed necessary for maintenance of both *in vitro* and *in vivo* transformation phenotypes [17, 19, 21].

Rather than displaying neofunctionalization in EwS, our data suggest that NR0B1 fulfills its well documented role as a nuclear receptor co-regulator [24, 26, 36, 37]. We might hypothesize that previous observations of direct physical and functional interaction between NR0B1 and EWS::FLI1 are in part reflective of the widespread interaction between ETS factors and nuclear receptors [46–48], mediated in part through conserved LXXLL-like peptide motifs in the ETS-domain that are retained in the EWS::FLI1 fusion [49, 50]. Therefore, EWS::FLI1 interactions with NR0B1 may involve their mutual association with nuclear receptors, and further investigation of this potential gene regulatory network is merited.

While we found distinct genome-wide binding patterns for NR0B1 across multiple EwS cell lines and PDX tissues, we observed highly convergent biological processes regulated by NR0B1 in each model system. We highlight NR0B1 activation of neural signatures including components of the npBAF chromatin remodeling complex and repression of innate immune and hormone response pathways. At the time this preprint was prepared, a zebrafish model demonstrating neural crest origins for EwS has been reported [51], consistent with previous findings that human neural crest stem cells tolerate EWS::FLI1 expression, resulting in loss of cellular senescence [52]. There are case reports of EwS tumors with histologic neural features [53], and previous studies showed that EwS cells have the plasticity to undergo neuronal differentiation upon loss of CD99 [54, 55]. Our findings suggest a possible role for NR0B1 in reinforcing signatures of partial neural differentiation in EwS, whether inherited from the cell of origin or established during tumor development. Dampening of hormone signaling programs by NR0B1 is well established in a variety of settings [24], yet here we find that NR0B1 may also have a direct role in suppressing EwS cell-intrinsic Type I interferon signaling. In particular, we found that NR0B1 loss enhanced the expression of *CGAS, IRF9/7/1, STAT1/2,* and *IFIT1/2/3/5*. While the impact of interferon signaling on cancer growth and metastasis appears complex and context specific in EwS and in cancer more broadly [56–59], it may be of value to dissect the cell autonomous effects of NR0B1 innate immune modulatory behavior as well as the broader impact on the tumor microenvironment in EwS [60].

With the notable exceptions of neuroblastoma and acute promyelocytic leukemia (APL), investigations of nuclear receptors as important mediators of pediatric cancer outcomes have been limited. This large family of tissue-specific transcription factors controls myriad metabolic, homeostatic, and developmental processes often in response to ligand stimulation [61]. They are therefore an attractive target for therapeutic intervention. Retinoids, signaling through RXR and RAR nuclear receptor subfamily members, emerged as indispensable treatments in neuroblastoma and APL by inducing terminal cellular differentiation, and there has been concerted effort to expand this success to additional cancer settings [62]. In osteosarcoma cells, retinoids, as well as ligands for additional nuclear receptors (e.g., PPARg), appear to effectively inhibit cancer cell proliferation *in vitro* and enhance the *in vivo* effects of doxorubicin [63], and in rhabdomyosarcoma cells, retinoids induce muscle differentiation [64]. While data is conflicting in EwS, previous studies showed that EwS cells were unresponsive to retinoids [65]. Further, activation of EWS::FLI1 (and presumably NR0B1) in embryonal carcinoma cells impeded their capacity for neuronal differentiation in response to retinoids [66]. Still a recent study found that SK-ES-1 EwS cells have reduced cell proliferation and spheroid formation capacity when exposed to high levels of retinoic acid [67], illuminating the potential for therapeutic retinoid responses in this disease. Our data demonstrate a strong physical and functional interplay between NR0B1 and RXRa in EwS cells. In concordance with extensive literature precedent that NR0B1 disrupts the ligand-dependent gene regulatory function of numerous nuclear receptors, we find evidence that NR0B1 expression and activity have dramatic impacts on the EwS response to 9cRA. Consequently, disruption of NR0B1 opens the door to a more robust retinoid response, activating adipogenic and myogenic differentiation pathways and repressing MYC targets as well as gene signatures characteristic of EwS. Our finding that ATRA (RAR ligand) fails to phenocopy 9cRA (pan-RAR/RXR ligand) treatment suggests that RXRs play a key role in mounting this response, and future efforts should endeavor to dissect the value of individual retinoid receptors as targets in EwS.

Finally, we highlight the recent development of BPK-29, an NR0B1 inhibitor that has afforded us the ability to investigate NR0B1 activity pharmacologically. *KEAP1*-mutant non small-cell lung cancer (NSCLC) is another rare example of aberrant NR0B1 transcriptional activation in cancer [68]. In this setting, Bar-Peled et al., identified Cysteine 274 of NR0B1 as a target for covalent inhibition and developed BPK-29, which disrupts interactions between NR0B1 and the nuclear receptor coactivator SNW1, as well as inhibits the anchorage-independent growth of *KEAP1-*mutant NSCLC cells (at 5 μM concentration) [28]. In our hands, BPK-29 treatment by itself had minimal impact on the gene expression profile of EwS cells but faithfully reproduced the effect of NR0B1 silencing with respect to 9cRA-dependent gene expression changes at 50-fold lower concentration than previously utilized. Our work reveals the tremendous potential of this compound to relieve cancer cells of the detrimental impacts of NR0B1 activity on the function of their endogenous nuclear receptors, potentially paving the way for a therapeutic response to individual or combinatorial nuclear receptor stimulation. Our early results suggest some interaction between BPK-29 and 9cRA in cell proliferation and transformation assays, encouraging further investigation of this or related treatment strategies. Our phenotypic analyses should be considered in the context of the marked intra- and intertumoral heterogeneity among EwS cells, which was reviewed recently [69]. We feel the phenotypic outcomes of NR0B1 disruption in combination with 9cRA stand alone as they demonstrate maintenance of high EWS::FLI1 transcriptional activity combined with increased 2D cell proliferation and reduced transformation. By largely disentangling EWS::FLI1 and NR0B1 activity, we are optimistic that ongoing work will allow us to better characterize this emergent EwS cell state and its sensitivity to new treatments.

## Acknowledgements

We would like to acknowledge the Institute for Genomic Medicine Genomic Services Lab for assisting with RNA-seq, ATAC-seq, and 10X snRNA/ATAC-seq data generation and the Institute for Genomic Medicine Computational Genomics Group for assistance in data curation and analysis. We are grateful to Dr. Katherine Miller and Dr. Genevieve Kendall for helpful discussion and input on the project. RDD is supported by a Presidential Fellowship from the Ohio State University. Funding was provided by the CancerFree KIDS NEW Idea Award (to BDS), Ohio Lion’s Pediatric Cancer Foundation/Nationwide Children’s Hospital Foundation (NCHF) (to BDS), Nationwide Children’s Hospital Center for Childhood Cancer Research (to RDR), Germain Family Accelerator Fund/NCHF (to RDR), St. Baldrick’s Scholar Awad (to ERT), ACS Research Scholar Grant (RSG-22-118-01-DMC, to ERT), Community Choice Foundation/NCHF (to ERM), and Nationwide Children’s Hospital Institute for Genomic Medicine/NCHF (to RKW).

## Author Contributions

Conceptualization – BDS

Methodology – BDS, RDD, MS, LR, ACG

Formal Analysis – BDS, MW, SF

Data Curation – BDS, MW

Resources – BDS, RDR, ERT, ERM, RKW

Writing – original draft: BDS – reviewing and editing: BDS, ERT, ERM

Supervision – BDS, RDR, ERT, ERM, RKW

Funding Acquisition – BDS, RDR, ERT, ERM, RKW

## Data and resource availability

All next-generation sequencing data will be uploaded to GEO/SRA and made publicly available upon final acceptance of this manuscript. No new code or resources were generated in the course of this work.

## Methods

### Cell culture

A673 (ATCC) and SKNMC (ATCC) cells were maintained in DMEM (Gibco 10-313-039) supplemented with 10% FBS, 1X Glutamax, and 1X Penicillin/Streptomycin. TC32 and TC71 cells were maintained in RPMI 1640 (Gibco 21-870-092) supplemented with 10% FBS, 1X Glutamax, and 1X Penicillin/Streptomycin. For siRNA transfections, cells were seeded one day prior to transfection and allowed to attach. Prior to transfection, media was replaced with fresh media with no antibiotics. siRNAs were prepared in Opti-MEM (Gibco 31-985-070) with Lipofectamine 3000 reagent. siRNA solutions were added dropwise to a final concentration of 67 μM and cells were incubated for 72 hours prior to collection. 9-*cis*-retinoic acid (Sigma R4643-1MG), all-*trans-*retinoic acid (Sigma R2625-50MG), and BPK-29 (Sigma SML2305-5MG) were prepared in DMSO. For ChIP- and RNA-seq assays, all compound treatments were at 100 nM for 24 hours.

### Western blot

50 mg of PDX tissue was minced in ice cold PBS, pelleted at 500 x g for 5 minutes at 4°C, and resuspended in RIPA buffer (200 mM NaCl, 50 mM Tris-HCl pH 8, 0.5% NP-40, 0.5% sodium deoxycholate, and 0.1% SDS) supplemented with protease inhibitors (Pierce PI78429). Tissue was transferred to a Covaris AFA milliTUBE and sonicated on an ME220 instrument. Remaining tissue debris was pelleted by centrifugation at 21,000 x g for 10 minutes at 4°C, and lysates were quantified by BCA assay (Pierce A55861). 10 μg of protein for each sample was resolved on a 4-12% Bis-Tris gel (NuPAGE) for 90 minutes at 100 V. Proteins were transferred to a nitrocellulose membrane overnight at 16 V. Membranes were blocked with 5% milk powder in TBST (0.1% Triton X100) before primary antibody incubation (NR0B1 – Cell Signaling Technology 13538S or ACTB – Cell Signaling Technology 3700S) at room temperature for 2 hours. After secondary antibody incubation for 1 hour at room temperature, membranes were imaged on a ChemiDoc imagine system with SuperSignal West PICO Plus Chemiluminescent Substrate.

### ChIP-seq and analysis

Chromatin was prepared using the Covaris TruChIP Chromatin Shearing Kit. Briefly, cells were typsinized and washed with PBS at room temperature. For NR0B1 ChIP, cells were first suspended in 2 mM disuccinimidyl glutarate (DSG, Pierce PIA35392) for 30 minutes, washed with PBS, and suspended in 1% formaldehyde for 10 minutes. Crosslinking was quenched with 125 mM glycine and fixed cells were washed twice with ice cold PBS, snap frozen, and stored until further use. For all other ChIP targets, we used single fixation with 1% formaldehyde.

TruChIP reagents were utilized to isolate nuclei from fixed cells, which were then suspended in TruChIP shearing buffer in a 1 mL AFA milliTUBE. Chromatin was sheared on a Covaris ME220 instrument and any remaining debris was pelleted by centrifugation at 21,000 x g for 10 minutes at 4°C. 20 uL of sheared chromatin was set aside as input (diluted to 100 μL in TE with 2.5 μL of 10% SDS and 5 μL of Proteinase K [Thermo EO0491]), and chromatin was diluted with 2 volumes of ChIP Buffer (0.1% SDS, 0.1% sodium deoxycholate, 1% Triton X100, 200 mM NaCl in 1X TE buffer). ChIP antibodies were then added, and the chromatin sample was rotated at 4°C for 1 hour prior to the addition of 50 μL of Protein A Dynabeads (Thermo 10001D) and further incubation overnight. Beads were washed twice with ice cold No Salt ChIP Buffer (0.1% SDS, 0.1% sodium deoxycholate, 1% Triton X100 in 1X TE buffer), ChIP Buffer, LiCl Buffer (0.5% NP-40, 0.5% sodium deoxycholate, 250 mM LiCl, in 1X TE buffer), and 1X TE buffer and finally suspended in 100 μL TE with 2.5 μL of 10% SDS and 5 μL of Proteinase K. ChIP and Input samples were decrosslinked overnight at 65°C and DNA was isolated with a QIAquick PCR Purification kit. ChIP and Input samples were quantified by Qubit, and Tapestation was used to confirm shearing efficiency.

ChIP-seq libraries were prepared using the NEBNext Ultra II DNA Library Prep Kit in combination with Singular Genomics Universal Library Prep Adapter and Unique Dual Index Primers. Libraries were sequenced on the Singular Genomics G4 instrument using F3 100-cycle reagent kits. Fastq files from the G4 were analyzed with the Encode ChIP-seq Pipeline 2 (v1.3.4), aligning to hg38. Comparison of NR0B1 binding sites across tissues was performed with the chipseeker, and all other peak (.bed) file manipulations were performed with bedtools. Heatmaps and profile plots were generated with deeptools, and motif analyes were performed with Homer. Chromatin states were determined using ChromHMM.

### RNA-seq and analysis

RNA samples were collected using the Qiangen QIAshredder and RNeasy Mini Kits with on-column DNase digestion. RNA samples were quantified by Nanodrop and submitted to the Genomic Services Laboratory in the Institute for Genomic Medicine (IGM) at Nationwide Children’s Hospital (NCH). Following determination of RNA quality (RIN) by Tapestation, RNA-seq libraries were prepared using the NEBNext Ultra II Directional RNA Library Prep Kit for Illumina. Pooled libraries were sequenced on the NovaSeq 6000 platform, and fastqs were analyzed using the nf-core RNA-seq pipeline aligning to hg38 (star_salmon option). Differential expression was performed using DESEQ2. Pathway analyses were performed by combining DESEQ2 differential gene expression profiles with ChIP-seq peak calls (e.g., NR0B1 peaks and NR0B1-dependent gene expression changes) on the Cistrome-GO platform.

### ATAC-seq and analysis

ATAC-seq was performed on 50,000 cells using the Active Motif ATAC-seq Kit per the manufacturer’s protocol. Pooled libraries were submitted to the Genomic Services Lab at IGM and sequenced on the NovaSeq 6000 platform. Fastqs were analyzed using the nf-core ATAC-seq pipeline, aligning to hg38. Differential peaks were identified using the DiffBind package. ATAC-seq heatmaps were prepared with deeptools and motif analysis was performed for Homer.

### PDX single nuclei RNA/ATAC-seq and analysis

PDX tissues (NCH-EWS-3 and NCH-EWS-5) were propagated and collected in the Animal Tumor Core at NCH with approval from the Institutional Animal Care and Use Committee (IACUC protocol AR15-00022). Tissues were lysed and nuclei were isolated and quantified using a hemocytometer. Nuclei were carried into the Chromium Next GEM Single Cell Multiomic ATAC + Gene Expression kit and libraries were prepared according to the manufacturer’s protocol. Gene expression and ATAC libraries were sequenced separately on the Illumina NovaSeq X Plus platform with a 10B 300 cycle reagent kit.

For analysis, we first prepared a joint hg38 and mm10 reference genome and transcriptome with cellranger arc (version 2.0.2). We then aligned the data to this joint reference genome/transcriptome. Seurat (version 5.1.0) and Signac (version 1.14.0) were used for QC and analysis. Nuclei with fewer than 1000 reads aligned to hg38 (for both ATAC and RNA) were discarded. Then, nCount and nFeature outliers for both RNA and ATAC assays were removed on a per sample basis. Nuclei with more than 5% mitochondrial RNA content in RNA assay were also removed. In the end there were 8278 nuclei for NCH-EWS-3 and 7266 nuclei for NCH-EWS-5. A weighted nearest neighbor network was built based on PCA (for RNA assay) and LSI (ATAC assay) with FindMultiModalNeighbors() function from Seurat. Standard clustering with FindClusters and UMAP projection were both constructed using the multimodal wsnn. Clustering resolution was chosen on a per sample basis through clustree visualization. Clusters were defined as “NR0B1 high” if NR0B1 expression was observed in more than 20% of nuclei within that cluster. Differential gene expression analysis was performed between the NR0B1 high and low groups. FindMarkers was used for the analysis, where logfc.threshold was set at 0.25 and min.pct at 0.1. ATAC peaks for NR0B1 high and low clusters were called with the CallPeaks() function from the signac package. Differential Accessibility analysis between the two clusters was performed with FindMarkers() function with likelihood ratio test option (test.use = ’LR’), latent.vars set as nCount_ATAC, and min.pct set at 0.1.

### Soft agar colony formation assay

Soft agar assays were performed as previously described [70]. For the A673 cell line, 7,500 cells were seeded per agar. Each plate was fed twice a week with 500 uL of media supplemented with drug and then imaged at 2 weeks post seed. Colony number, average colony size, and percent area were counted using ImageJ software.

### Cell proliferation and GILA assays

For 2D cell proliferation, EwS cell were plated at a density of 5,000 cells in 100 μL per well of a 96-well tissue culture plate and allowed to attach overnight. The following day (Day 0), cells were treated with all combinations of 9cRA (at 0, 1, 5, 10, 50, 100, 500, and 1000 nM) and BPK-29 (at 0, 10, 50, 100, 250, 500, 750, and 1000 nM). Media and compounds were replaced on Days 2 and 4. On Day 6, 0.1 volumes of PrestoBlue HS Cell Viability Reagent were added to each well, and the cells were returned to the incubator for 6 hours. Absorbance was measured at 570 nm and normalized to absorbance at 600 nm. % viability was calculated based on the corrected absorbance for each well with respect to DMSO treatment (i.e., 0 nM 9cRA and 0 nM BPK-29). Replicate results were analyzed with SynergyFinder to determine single agent IC50 values and calculate synergy/antagonism.

For Growth in Low Attachment (GILA) assays, 10,000 EwS cells in 100 μL were seeded in 96-well ultra-low attachment, flat bottom plates (Corning 3474). The following day (Day 0), cells were treated with all combinations of 9cRA (at 0, 1, 5, 10, 50, 100, 500, and 1000 nM) and BPK-29 (at 0, 10, 50, 100, 250, 500, 750, and 1000 nM). Compounds were refreshed without media change on Days 2 and 4. On Day 6, 0.1 volumes of PrestoBlue HS Cell Viability Reagent were added to each well, and the cells were returned to the incubator for 6 hours. Absorbance was measured at 570 nm and normalized to absorbance at 600 nm. Corrected absorbance values for each assay plate were normalized to be between 0 and 1, and % viability was calculated for each well with respect to DMSO treatment (i.e., 0 nM 9cRA and 0 nM BPK-29). Results were analyzed with SynergyFinder to calculate synergy/antagonism.

## Materials

### Antibodies

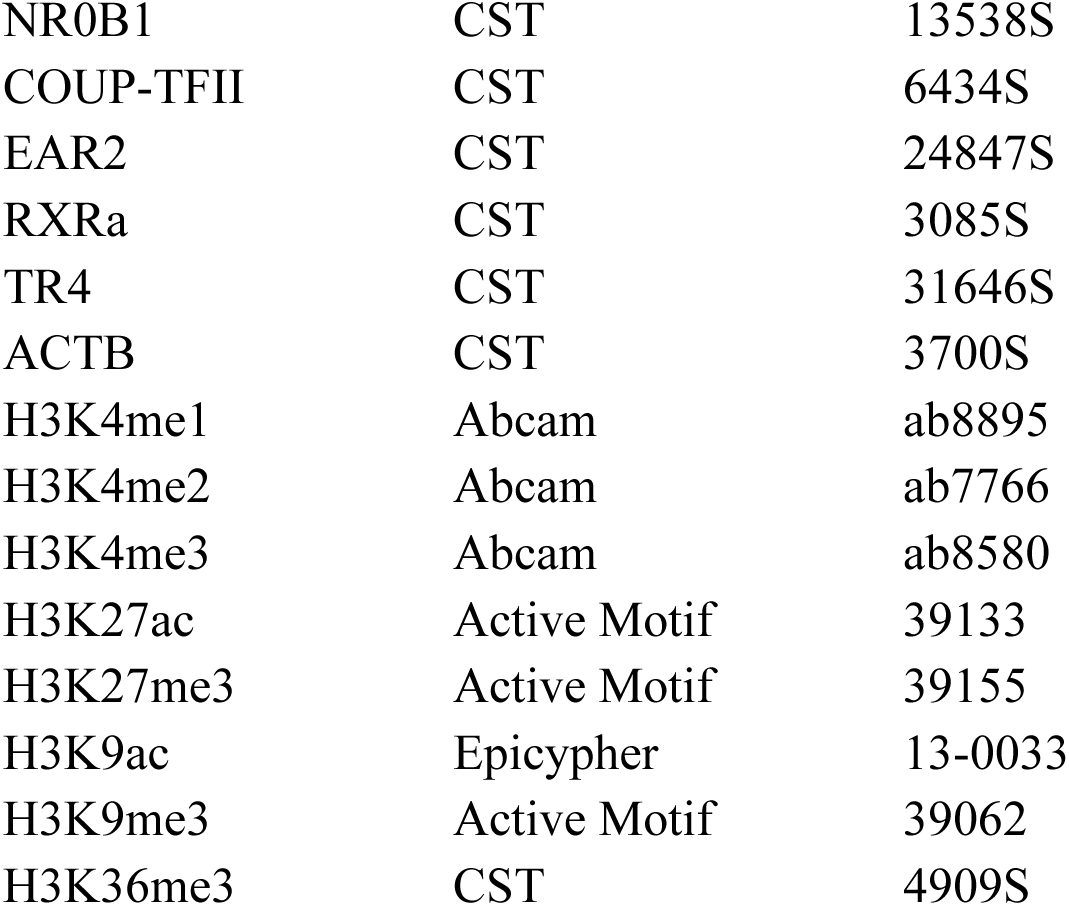

### Public Data

|  |  |  |
| --- | --- | --- |
| EWS::FLI1 ChIP-seq | GSE92741 | [35] |
| Tumor Expression Data | St. Jude Cloud/PeCan | [71, 72] |
| EWS Survival Data | R2 Platform | [32, 33] |

---

**Supplementary Figure S1.**
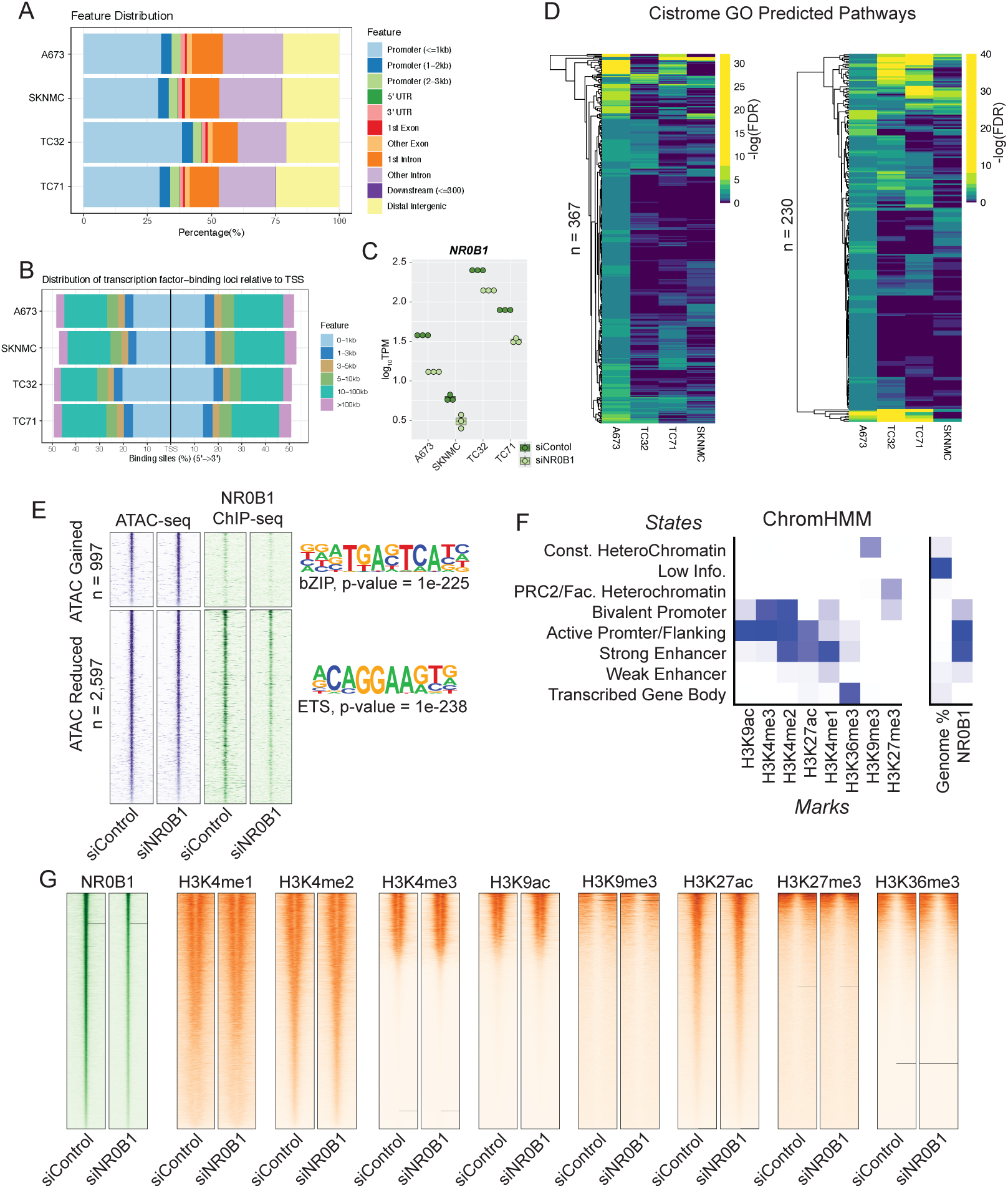
**A)** Chipseeker annotation of NR0B1 binding sites in the indicated genomic regions for A673, SKNMC, TC32, and TC71 cell lines. **B)** Chipseeker annotation of NR0B1 binding sites distances from all transcription start site (TSS) locations in the indicated cell lines. **C)** *NR0B1* TPM values from RNA-seq analysis in the indicated cell lines transfect with siControl or siNR0B1. **D)** Comparison of NR0B1 target gene pathway enrichment across the indicated cell lines for the 367 NR0B1 repressed and 230 NR0B1 activated pathways identified in A673 cells. **E)** Heatmap of ATAC-seq and NR0B1 ChIP-seq signal strength in regions of gained or reduced chromatin accessibility in A673 cells transfected with siControl vs. siNR0B1. Top motifs enriched in gained and reduced sites are provided. **F)** An 8-state ChromHMM model was generated in A673 cells transfected with siControl or siNR0B1. Heatmaps display the enrichment of 8 individual histone modifications as well as NR0B1 in each ChromHMM state. **G)** Heatmaps display the ChIP-seq signal intensity for NR0B1 and 8 histone marks across NR0B1 peaks in A673 cells transfected with siControl or siNR0B1. Peaks are ordered independently by descending strength for each factor profiled.

**Supplementary Figure S2.**
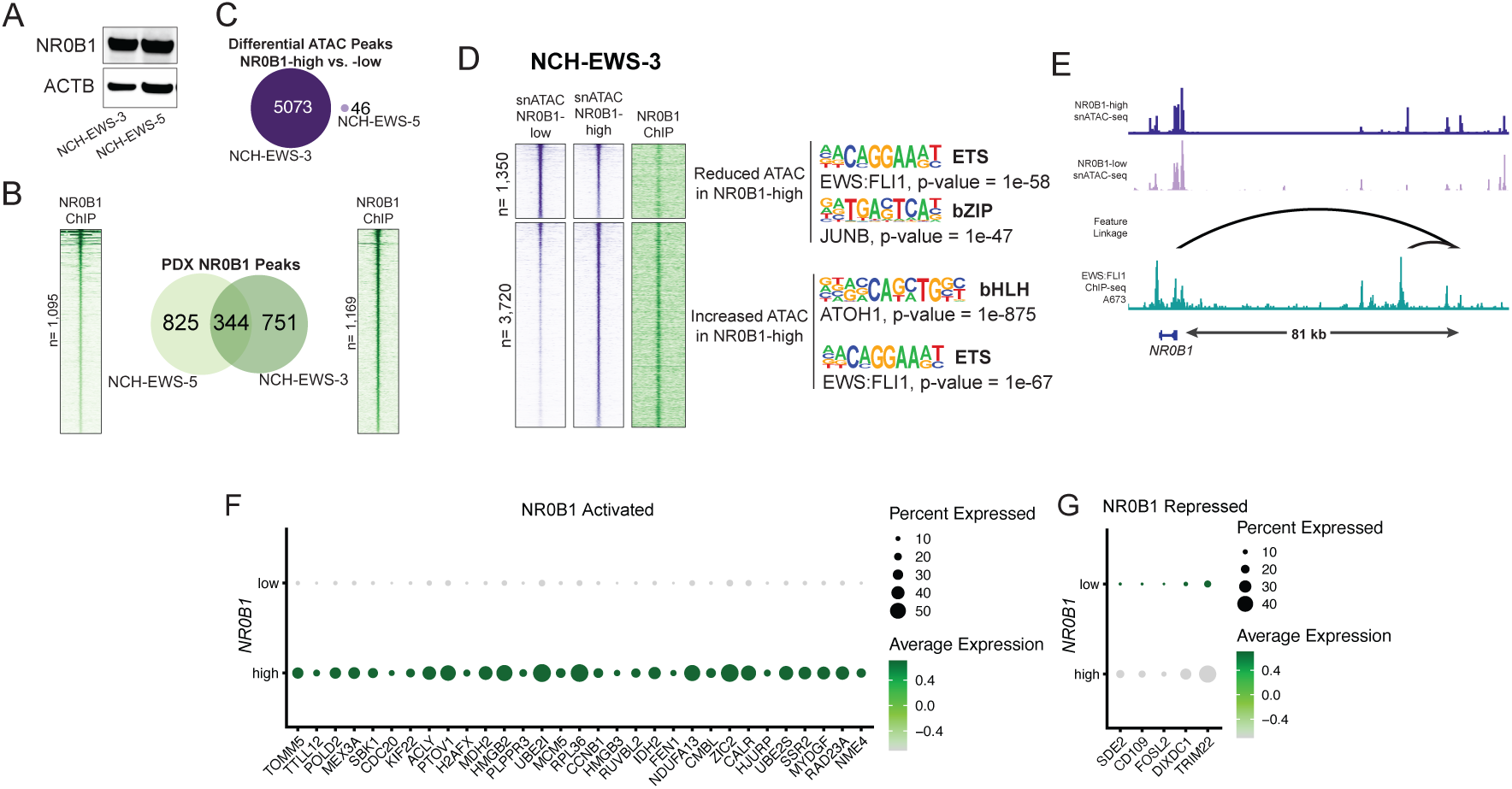
**A)** Western blot shows expression of NR0B1 (ACTB loading control) in NCH-EWS-3 and NCH-EWS-5 PDX tissues. **B)** Bulk NR0B1 ChIP-seq was performed in NCH-EWS-3 and NCH-EWS-5 PDX tissues. Heatmaps show NR0B1 ChIP-seq signal strength across all binding sites. Venn diagram displays unique and shared NR0B1 binding sites between PDX tissues. **C)** Differential accessibility was determined for the indicated PDX tissues, comparing NR0B1 high nuclei to NR0B1 low nuclei. **D)** Aggregate snATAC-seq signal intensity from NCH-EWS-3 NR0B1 high vs. low nuclei clusters displayed over differential ATAC peaks. Bulk NR0B1 ChIP-seq signal strength and enriched motifs are also provided. **E)** IGV plot of aggregate snATAC-seq signals from NR0B1 high vs. low nuclei clusters and feature links from NCH-EWS-3 at the *NR0B1* gene locus. EWS::FLI1 ChIP-seq data from A673 cells is also provided. **F-G)** Genes activated or repressed by NR0B1 in NCH-EWS-5 as well as in A673 cells were uploaded to the R2 platform to conduct Kaplan-Meier survival analysis. Dot plot depicts the expression patterns (in NR0B1 high vs. low nuclei) of NR0B1 activated genes for which high expression in EwS patients is significantly associated with poor overall survival **(F)** and NR0B1 repressed genes for which low expression is associated with poor overall survival **(G)**.

**Supplementary Figure S3.**
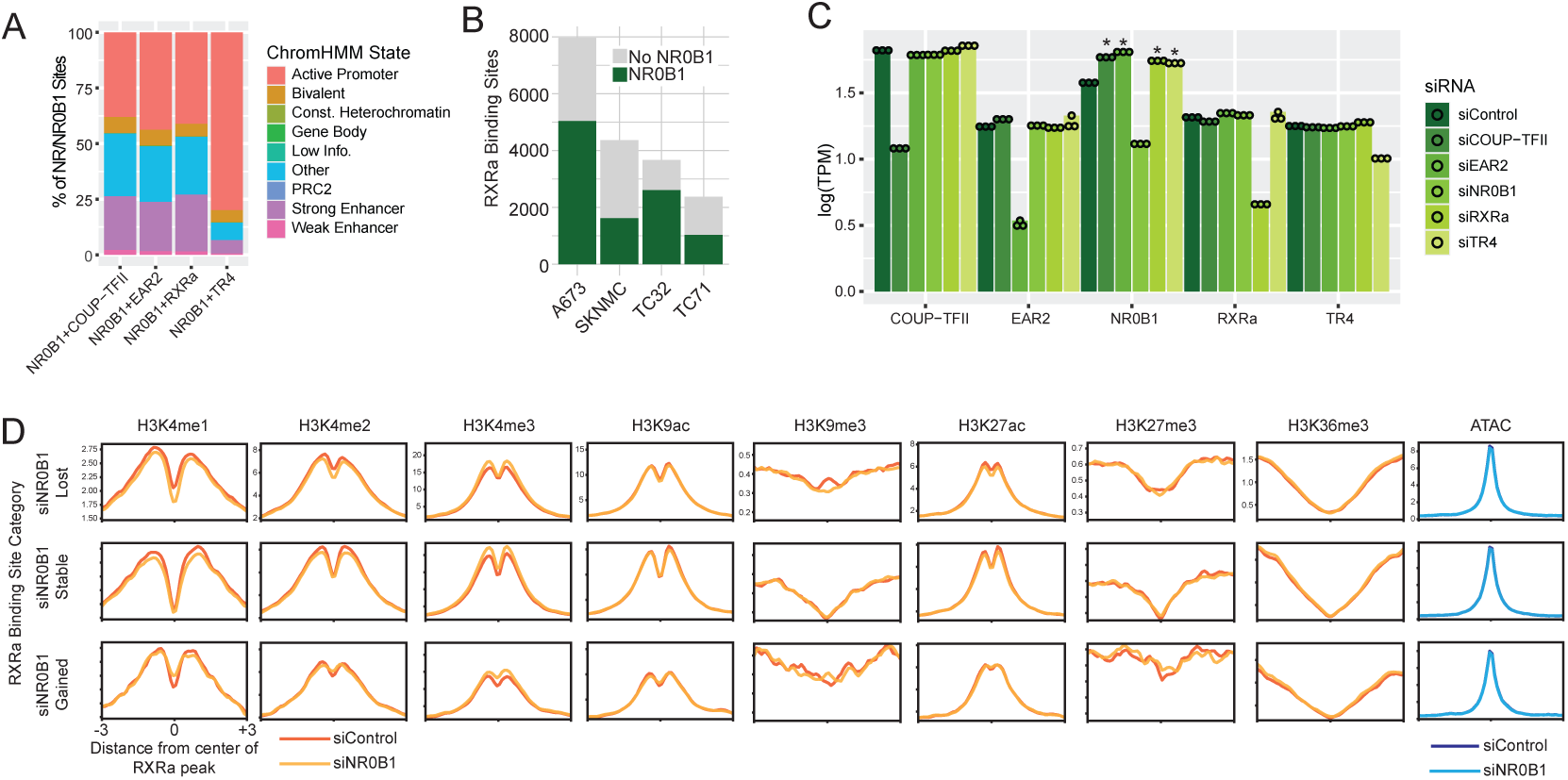
**A)** Overlap of sites co-occupied by NR0B1 and COUP-TFII, EAR2, RXRa, or TR4 with ChromHMM states in A673 cells. **B)** NR0B1 overlap with RXRa binding sites in the indicated EwS cell lines. **C)** log_10_(TPM) values for *COUP-TFII, EAR2, NR0B1, RXRa,* and *TR4* in A673 cells transfected with the indicated siRNA. * indicates significant differential expression compared to siControl. **D)** Aggregate ChIP-seq profile plots for the indicated epigenetic markers across RXRa sites lost, stable, or gained as a result of NR0B1 silencing

**Supplementary Figure S4.**
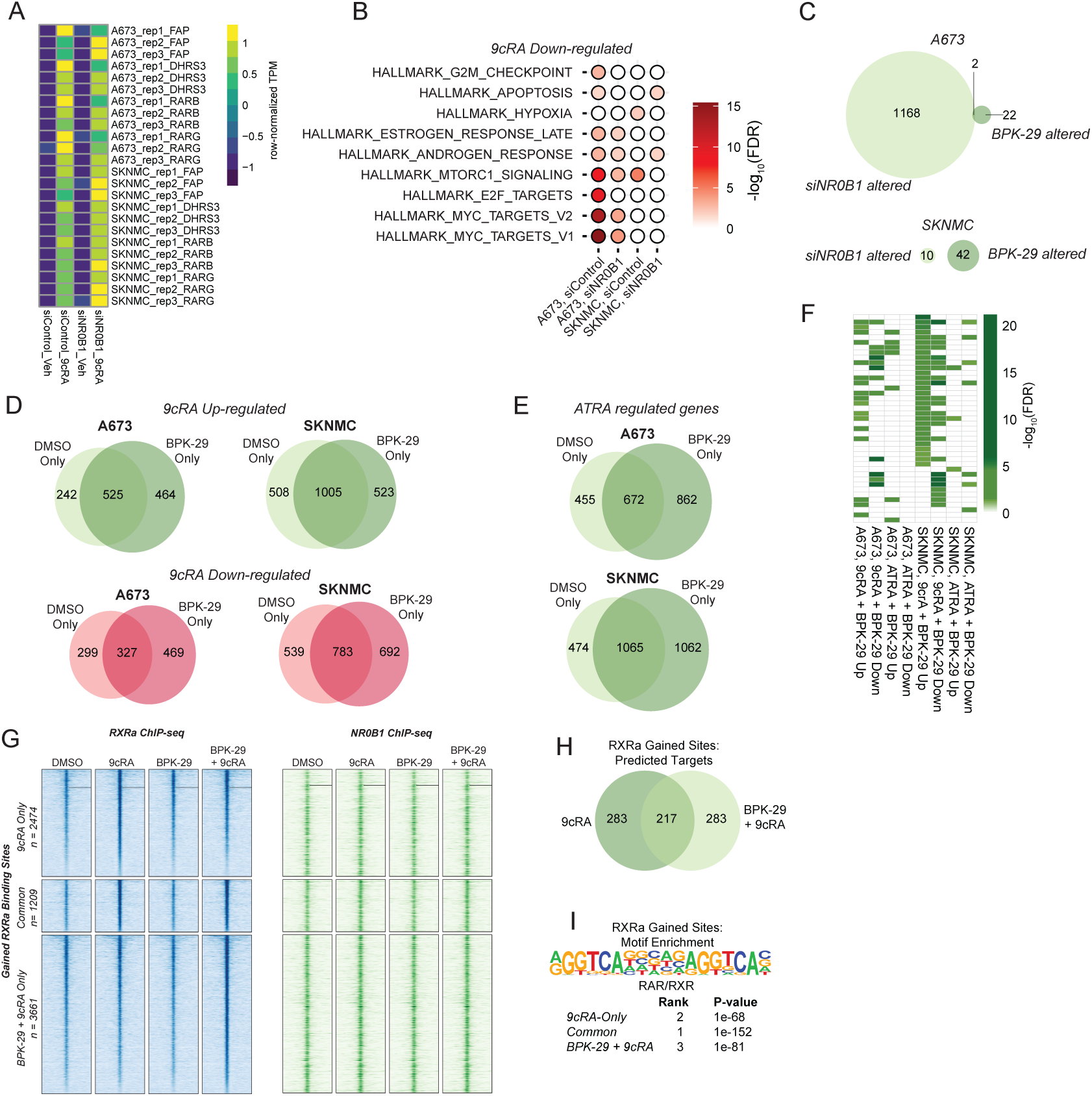
**A)** Relative expression level of canonical retinoid inducible genes (*FAP, DHRS3, RARB, RARG*) in A673 and SKNMC cells transfected with siControl or siNR0B1 and treated with Vehicle (DMSO) or 9cRA. **B)** Genes uniquely down-regulated by 9cRA in siControl vs. siNR0B1 transfected cells were used to assess enrichment of MSigDB Hallmark Pathways. Balloon plot depicts pathway enrichment in siControl vs. siNR0V1 transfected cells. **C)** Comparison of siNR0B1 altered vs. BPK-29 altered genes in A673 and SKNMC cells. **D)** RNA-seq was performed in A673 and SKNMC cells treated for 24 hours with 100 nM 9cRA with/without 100 nM BPK-29. Differential expression analysis identified 9cRA responsive gene expression changes in DMSO vs. BPK-29 treated cells. Venn diagrams display the genes uniquely up-regulated or down-regulated by 9cRA in the presence vs. absence of BPK-29. **E)** Venn diagrams display the genes uniquely regulated by ATRA in the presence vs. absence of BPK-29 in A673 and SKNMC cells. **F)** Comparison of significantly enriched MSigDB Hallmark Pathwyas among genes up- or down-regulated by 9cRA vs. ATRA in A673 cells cotreated with BPK-29. **G)** RXRa and NR0B1 ChIP-seq signal strength in RXRa binding sites enhanced by 9cRA alone, 9cRA + BPK-29 alone, or by either treatment condition. **H)** RXRa binding sites enhanced only by 9cRA or only by 9cRA + BPK-29 treatment were combined with 9cRA-driven gene expression changes (+/- BPK-29) on the Cistrome GO platform to predict putative RXRa target genes in each condition. Venn diagram compares the top 500 predicted RXRa target in each condition. **I)** Motif analysis of RXRa binding sites enhanced by 9cRA alone, 9cRA + BPK-29 alone, or by either treatment condition.

**Supplementary Figure S5.**
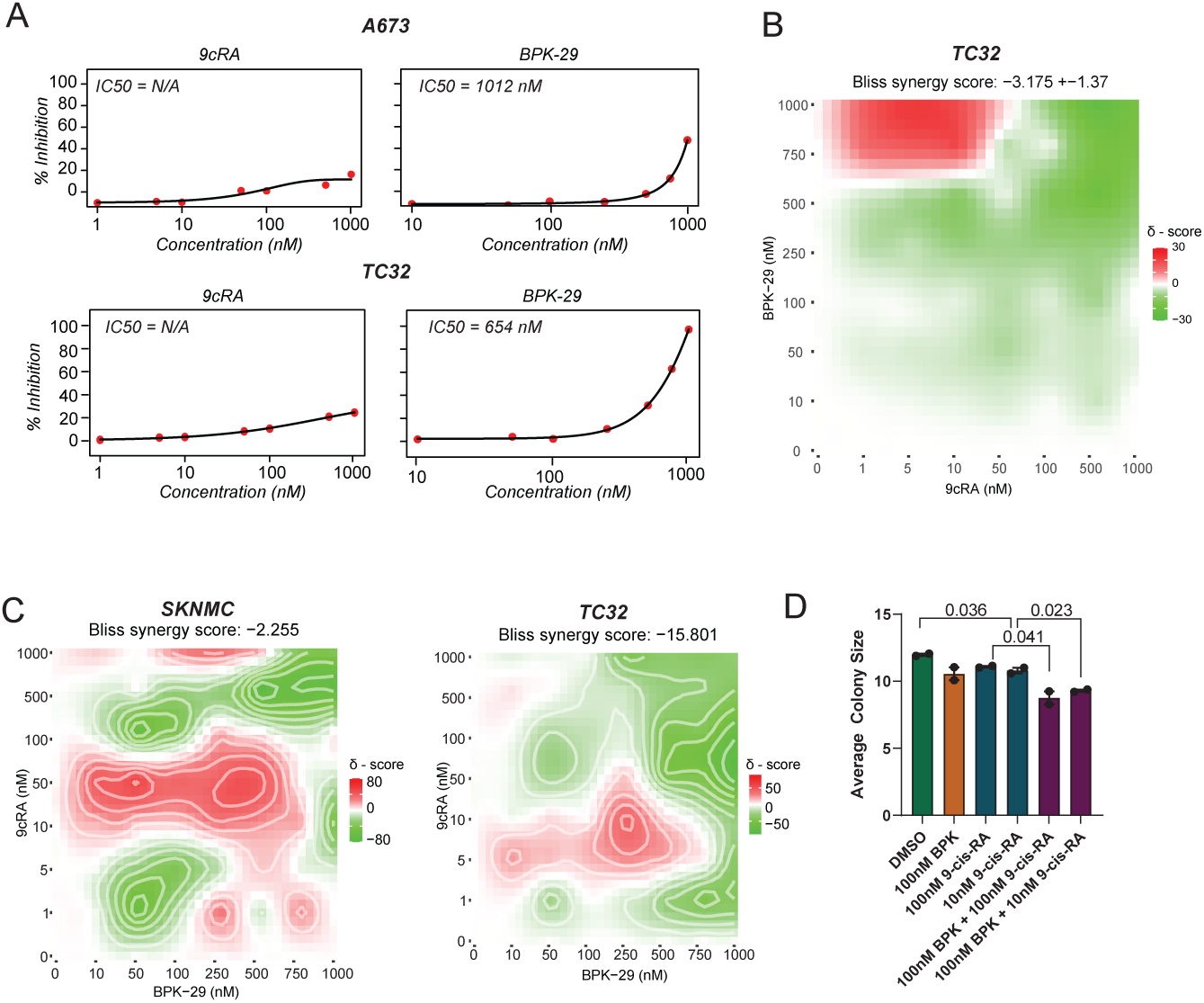
**A)** Dose-response curves for 9cRA or BPK-29 in A673 and SKNMC cells grown as a monolayer. **B)** TC32 cell grown in a monolayer were treated with the indicated combinations of 9cRA and BPK-29 for 6 days, and cell viability was measured. Results were analyzed on the SynergyFinder platform, demonstrating antagonism between 9cRA and BPK-29. **C)** SKNMC and TC32 spheroids were grown in ultra-low attachment plates exposed to the indicated combinations of 9cRA and BPK-29 for 6 days. Viability results were analyzed on the SynergyFinder platform, demonstrating compound interaction between 9cRA and BPK-29. **D)** Soft agar colony formation assays were performed on A673 cells exposed to the indicated treatments. Average colony sizes (pixels) are provided as well as p-values (t-test) for significant comparisons.

